# SummaryAUC: a tool for evaluating the performance of polygenic risk prediction models in validation datasets with only summary level statistics

**DOI:** 10.1101/359463

**Authors:** Lei Song, Aiyi Liu, Molecular Genetics of Schizophrenia Consortium, Jianxin Shi

## Abstract

**Motivation:** Polygenic risk score (PRS) methods based on genome-wide association studies (GWAS) have a potential for predicting the risk of developing complex diseases and are expected to become more accurate with larger training data sets and innovative statistical methods. The area under the ROC curve (AUC) is often used to evaluate the performance of PRSs, which requires individual genotypic and phenotypic data in an independent GWAS validation dataset. We are motivated to develop methods for approximating AUC of PRSs based on the summary level data of the validation dataset, which will greatly facilitate the development of PRS models for complex diseases.

**Results:** We develop statistical methods and an R package SummaryAUC for approximating the AUC and its variance of a PRS when only the summary level data of the validation dataset are available. SummaryAUC can be applied to PRSs with SNPs either genotyped or imputed in the validation dataset. We examined the performance of SummaryAUC using a large-scale GWAS of schizophrenia. SummaryAUC provides accurate approximations to AUCs and their variances. The bias of AUC is typically less than 0.5% in most analyses. SummaryAUC cannot be applied to PRSs that use all SNPs in the genome because it is computationally prohibitive.

**Availability:** https://github.com/lsncibb/SummaryAUC

## 1. INTRODUCTION

Large-scale genome-wide association studies (GWAS) have identified dozens or even hundreds of common SNPs associated with many complex diseases, including psychiatric conditions, e.g., Schizophrenia (Schizophrenia Working Group of the Psychiatric Genomics, 2014), type 2 diabetes (Scott, et al., 2017) and common cancers, e.g., breast cancer (Michailidou, et al., 2017) and prostate cancers (Al Olama, et al., 2014). Heritability analysis using algorithms such as GCTA (Yang, et al., 2010) and LD-score regressions (Bulik-Sullivan, et al., 2015) have shown that, for many complex diseases, common SNPs have the potential to explain substantially larger fraction of the phenotypic variance than that based on the established GWAS SNPs, suggesting a great promise for genetic risk prediction. In fact, polygenic risk scores (PRSs) have been proven useful for predicting complex disease risk and are expected to become more accurate with large training data sets (Chatterjee, et al., 2013; Dudbridge, 2013) and innovative statistical methods. For example, a PRS for Schizophrenia based on tens of thousands of SNPs achieves an impressive prediction accuracy with an area under the receiver operating characteristic curve (AUC) of 0.75 (Schizophrenia Working Group of the Psychiatric Genomics, 2014) with approximately 35,000 cases and 45,000 controls.

Much effort has been invested on developing PRS prediction models and on investigating the factors that determine the performance of PRSs. When raw genotypic and phenotypic data are available for the training dataset, machine learning algorithms (Kooperberg, et al., 2010; Wei, et al., 2009), and linear mixed models (Golan and Rosset, 2014; Maier, et al., 2015; Speed and Balding, 2014) can be used to develop PRSs. PRSs can also be constructed when only GWAS summary level data are available for training dataset by simple *p*-value thresholding (Purcell, et al., 2009) or by more sophisticated methods that model linkage disequilibrium (Vilhjalmsson, et al., 2015). We have recently extended the p-value thresholding method to further improve accuracy by accounting for winner’s curse and by incorporating functional annotation data (Shi, et al., 2016). Different aspects of building PRS are discussed in a recent review paper (Chatterjee, et al., 2016).

Receiver operating characteristic (ROC) curve is one of the most popular tools for characterizing and comparing the diagnostic accuracy of binary classifier systems such as the PRSs in the present context. Since its first appearance in the Second World War for detecting enemy objects in battlefields, the ROC curve analysis has found its place in many other. A few books provide comprehensive coverage of the topics, see, among others (Hanley and Mcneil, 1982; Krzanowski, 2009; Pepe, 2003; Zou, 2011). The ROC curve of a PRS is generated by plotting its true positive rate against its false positive rate at various threshold. The area under the ROC curve (AUC) provides a quantitative measure for the discrimination ability of a PRS (Hanley and Mcneil, 1982). AUC is the most frequently used quantitative measure for evaluating the discrimination performance of a PRS, although some concerns have been raised for using AUC as a criterion for model comparison and risk stratification (Katki and Schiffman, 2018). While AUC is defined as the area under the ROC curve, i.e., the integral of the curve, a more convenient mathematical expression of AUC is the probability that a randomly selected case has a larger PRS value than a randomly selected control., With this expression, one can show that an AUC estimator is closely related with the Mann–Whitney U statistic and the Wilcoxon rank test. Given PRS values for a set of cases and controls, one can easily calculate AUC using this approach and estimate the variance of the estimated AUC using bootstrap.

Calculating AUC for a PRS typically requires the individual level genotypic and phenotypic data in an independent GWAS validation dataset. One solution is to genotype a large set of subjects as a new validation GWAS dataset, which is financially expensive and time consuming. Another possibility is to request individual level data from existing large scale GWAS independent of the training GWAS, which is also time consuming and may turn out to be infeasible because of data sharing policies. Instead, requesting summary statistics (odds ratio, *p*-value and imputation quality for individual SNPs) from an independent validation GWAS consortium is much easier because such summary statistics are usually available online with open access. Thus, developing methods for evaluating the performance of PRSs based on the summary statistics of validation GWAS would substantially accelerate the assessment of PRSs for specific diseases and facilitate the development of more accurate PRSs.

In this manuscript, we develop a statistical method, termed as “SummaryAUC”, for approximating AUC and its variance for a given PRS based on summary statistics from an independent GWAS validation data set. Although SummaryAUC relies on the normality assumption of PRS, extensive simulation studies demonstrate that it is highly accurate under realistic situations with more than 5 SNPs. Furthermore, SummaryAUC is flexible for PRSs with independent SNPs or SNPs in weak linkage disequilibrium (LD) and for both genotyped and imputed SNPs in the validation dataset. Finally, we applied SummaryAUC to schizophrenia GWAS to demonstrate the validity of the methods. SummaryAUC is best used for PRSs with independent SNPs and for PRSs with less than 20,000 SNPs in weak LD for both accuracy and computational efficiency. SummaryAUC is not suitable for PRSs integrating all common SNPs in the genome, e.g., LD-Pred (Vilhjalmsson, et al., 2015) and BLUP-type PRSs (Golan and Rosset, 2014; Speed and Balding, 2014) based on linear mixed models. An R package with the same name was developed to implement the proposed method and is publicly available.

## 2. METHODS

We assume an additive polygenic risk score (PRS) model based on *M* SNPs:

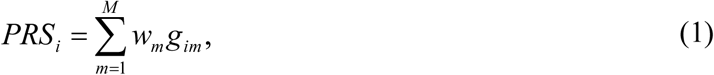

where *m* indexes SNPs and *i* indexes subjects in the validation data set. The weights (*w_l_*,· · ·, *w_M_*) are derived based on a specific algorithm and a training data set. The genotypic value *g_im_* ∈ {0,1,2} if the SNP is genotyped and *g_im_* ε [0,2] if the SNP is imputed. The selected SNPs in the PRS may be correlated because of linkage disequilibrium (LD).

When the genotypic data and the binary phenotypic data (*y_i_*) are available for each individual subject in the validation dataset, one can calculate PRS in (1) for all subjects and evaluate the performance of the prediction model by comparing PRS with the known phenotypic data. The performance of a prediction model is often assessed by the area under the receiver operating characteristic curve (AUC) at the observational scale.

We are interested in developing methods for estimating AUC and its standard deviations when only the GWAS summary statistics are available for the validation data set. For the *m*^th^ SNP, the summary statistics include the minor allele, the minor allele frequency (MAF) in the control samples, the odds ratio (*OR_m_*) or equivalently the regression coefficient *ß_m_* = log(*OR_m_*), the two-sided *P*-value *P_m_* or equivalently the *Z*-score statistic *Z_m_* = *sign* (*ß_m_*)Φ^−1^(1 - *P_m_*/2), the imputation information score 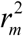, the number of cases *n*_1_ and the number of controls *n*_0_. Here, *ß_m_* and *P_m_* are based on single variant logistic regression. Φ() is the cumulative distribution function for *N*(0,1). In addition, MAF may not be available from the summary statistics to prevent subjects in the study to be deidentified (Homer, et al., 2008; Jacobs, et al., 2009).

### 2.1 Estimating AUC and its variance using summary statistics

Let 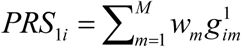 be PRS for the *i^th^* case and 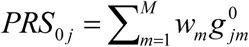 for the *j^th^* control subject. We assume that *M* is reasonably large that PRSs approximately follow normal distributions in cases and controls, respectively:

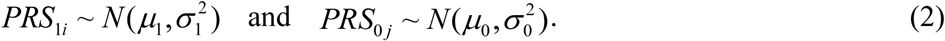

We will investigate the impact of the normality assumption in numerical studies. We define

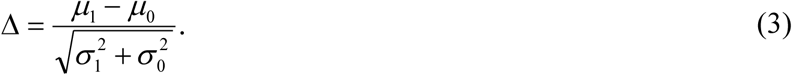

The AUC (denoted as *θ*) is defined as the probability that, for a randomly selected case and a randomly selected control, the case has a larger PRS than the control, i.e.,

*θ* = *P*(*PRS*_1*i*_ > *PRS*_0*j*_). For a validation data set with *n*_1_ cases and *n*_0_ controls with individual PRS values, one can estimate the AUC based on the following U-statistic (or the Wilcoxon-Mann-Whitney statistic):

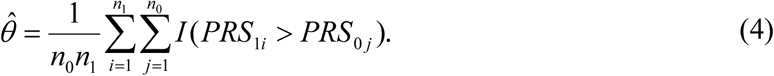

The expectation of AUC assuming normal distributions defined in (2) can be calculated as

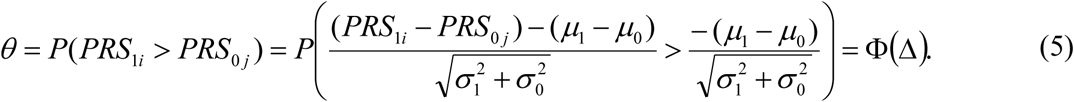

When *n* >> 1, *n*_0_ >> 1 and 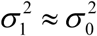 (when SNPs have modest effect sizes for complex diseases), we derive in Appendix that

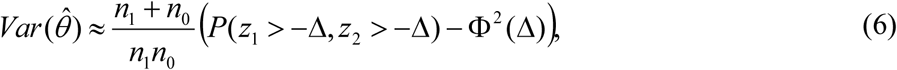

where *z*_1_~ *N*(0,1), *z*_2_~ *N*(0,1) and cor(*z*_1_,*z*_2_) = 1/2.

Note that both AUC (5) and the variance of the AUC estimator (6) depend on Δ defined in (3). Thus, it remains to estimate Δ based on the summary statistics in the validation dataset.

### 2.2 Estimate Δ when SNPs are independent

Let *p*_1*m*_ and *p*_0*m*_ denote the MAF of SNP *m* in the case and the control group, respectively. Typically, *p*_0*m*_ is included in the GWAS summary data while *p*_l *m*_ is not included. Let 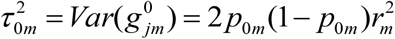 be the genotypic variance in the control group assuming the Hardy Weinberg Equilibrium law. Similarly, let 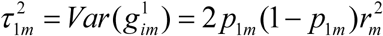 be the genotypic variance in the case group. Remember that we assume 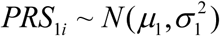 and 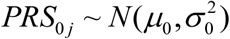 approximately in (2). When SNPs in the PRS are independent, we have

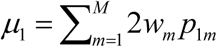, 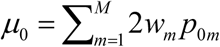, 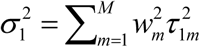, and 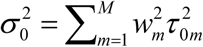 these into (3) leads to

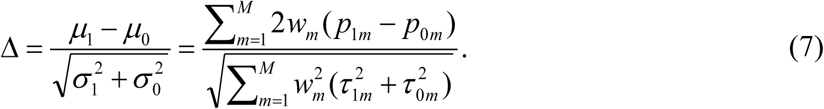

Thus, it remains to estimate *p*_1*m*_ – *p*_0*m*_, the difference of the allele frequencies between cases and controls.

Let 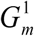 and 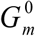 be the average genotypic values in the case group and the control group, respectively. One can estimate *p*_1*m*_ – *p*_0*m*_ as

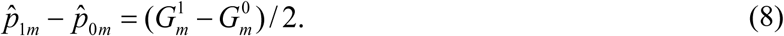

The *Z*-statistic for genetic association can be approximated by the *t*-statistic for large studies, *i.e.,* 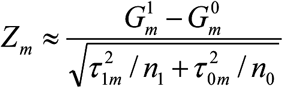 thus, we have

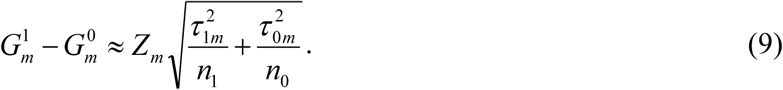

Combining (7), (8) and (9) leads to an estimate for Δ:

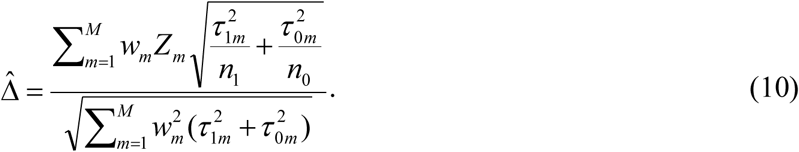

For complex diseases, 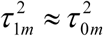 because common SNPs typically have small effects. Thus, we can use *p*_0*m*_ to calculate the genotypic variance 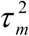 for both groups. When *p*_0*m*_ are not included in the summary statistics, we can use the allele frequency based on public data (e.g., The 1000 Genome Project) of the similar ancestry populations to approximate 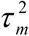. Under this assumption, (10) is simplified as

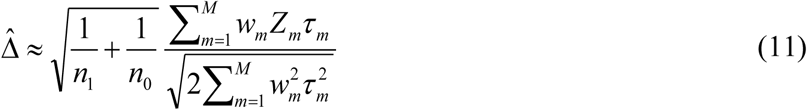

### 2.3 Estimate Δ when some SNPs are in linkage disequilibrium

We first assume that SNPs in PRS are genotyped in the validation data set. When some SNPs in PRS are in LD, we only need to modify the denominator in (11). We assume that, for complex diseases, correlations between local SNPs are similar in cases and controls. This assumption is reasonable because odds ratios are modest for nearly all disease-associated SNPs. Let *p*_*ml*_ = *cor*(*g_im_,g_il_*) be the genotypic correlation between SNP *m* and SNP *l*. The variance of PRS in controls and cases are 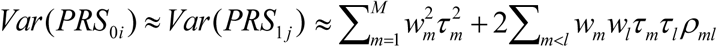. Thus

(11) can be modified as

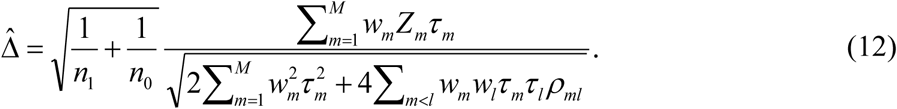

In implementation, we assume *p*_*ml*_ = 0 for SNPs located on different chromosomes and for SNPs on the same chromosome but more than 5 Mb away. In addition, we estimated *p_ml_* using the genotype data of the similar ancestry population in The 1000 Genomes Project.

However, it is very challenging when SNPs in the PRS are imputed. Let 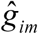 be the imputed genotypic dosage. Let 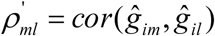 be the correlation between the imputed genotypic dosages. Although we cannot rigorously prove, we observe that imputation tends to inflate the magnitude of pairwise correlation, particularly for SNPs imputed with high uncertainty. We find that using *ρ*_*ml*_ (calculated based on genotype data in external data) to calculate 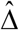 in (12) makes the approximation to AUC and its variance less accurate, particularly when a PRS has many SNPs. To address this problem, we propose a strategy to estimate 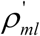 using The 1000 Genome Project data, which is illustrated in Figure 1.

**Figure 1.**
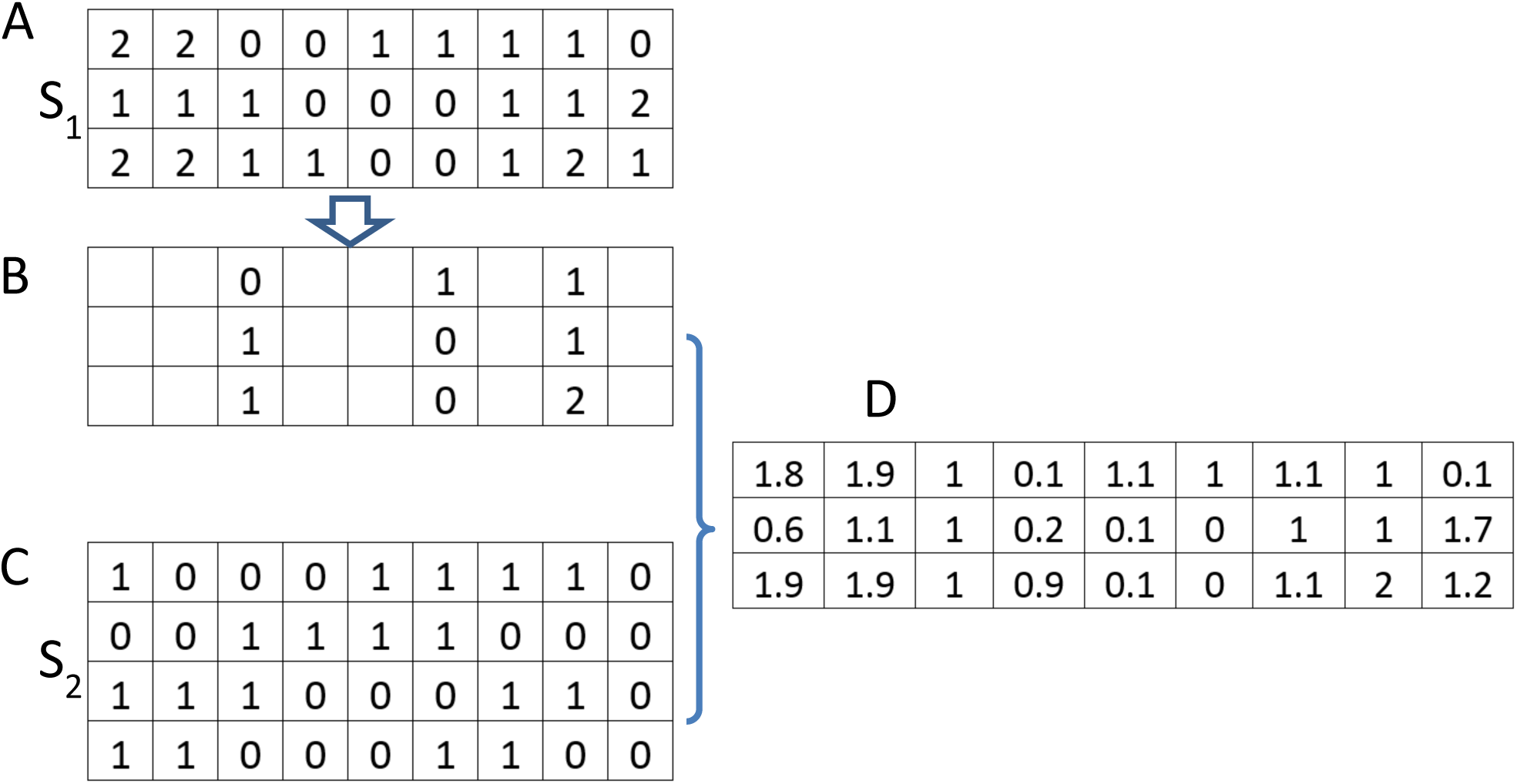
Estimate correlation of imputed SNPs in the validation GWAS dataset. The subjects in The 1000 Genome Project with relevant ancestry is divided as two sets S1 and S2. A: The genotype of the subjects in S1. B: Only SNPs that are genotyped in the validation GWAS dataset are kept. C: The haplotypes in S2 are used as reference panel for imputation. D. Imputation is performed to derive the genotypic dosage for SNPs that are not genotyped in the validation GWAS dataset. The correlation between two SNPs are calculated based on the imputed genotypic dosages.

Briefly, the subjects in The 1000 Genome Project with relevant ancestry are divided into two sets, denoted as S_1_ and S_2_ For subjects in S_1_, we keep only SNPs that are genotyped in the validation GWAS data set and perform imputation to derive genotypic dosages using the haplotypes in S_2_ as the reference panel. We can calculate 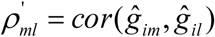 using the imputed genotypic dosages to approximate 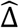 in (12). Obviously, the degree of uncertainty of imputed genotypes is similar to that in the validation GWAS data set because they are based on the same set of genotyped SNPs. Thus, we expect this strategy to attenuate the impact of imputation to the calculation of LD and thus to improve the approximation to AUC and its variance.

## 3. RESULTS

### 3.1 Implementation

We implemented our algorithms in an R package “SummaryAUC”, which is freely available online. Pairwise correlations between SNPs are estimated using the genotype data in the The 1000 Genome Project with relevant ancestry. For PRS with practically independent SNPs (e.g., SNPs pruned rigorously), there is no limitation on the number of SNPs in PRS. When SNPs in a PRS are correlated, we only adjust for correlations for SNP pairs that are less than 5 Mb away. In real data analyses, most PRSs have less than 10,000 SNPs, which can be calculated in a few minutes.

### 3.2 Simulation studies

The key assumption of SummaryAUC is that PRS follows a normal distribution approximately. This assumption may lead to poor approximation to AUC when the number of SNPs (*M*) in a PRS is small. Thus, we performed simulations to investigate whether and how the accuracy of SummaryAUC depends on *M*. In each simulation, we simulated genotypes for 3000 cases and 3000 controls and for *Μ* independent SNPs (*M* = 5,10, ···, 100). The allele frequency *p*_0*m*_ in controls followed a uniform distribution *U* (0.05,0.5). We simulated *ß*_*m*_ = *log (OR_m_*) *~N*(0,1/3^2^). The coefficients for PRS were set as the simulated *β* values.

For each set of simulated data, we calculated AUC and its variance in two ways. In the first approach, we calculated PRS for each individual subject using the individual genotypic values; we then calculated AUC using an R package “AUC” and its variance using bootstrap (*N* =10000). In the second approach, we first performed association test for each SNP to derive *Z*_*m*_ and also allele frequency *p*_0*m*_ for control samples; we then approximated AUC and its variance using SummaryAUC.

The simulation results are summarized in Figure 2. Results suggest that SummaryAUC provides very accurate approximation to AUC and its variance even when PRS has only five SNPs. Note that the correlation between true AUC and approximated AUC is greater than 99%. Thus, the performance of SummaryAUC is robust to the number of SNPs in PRS.

**Figure 2.**
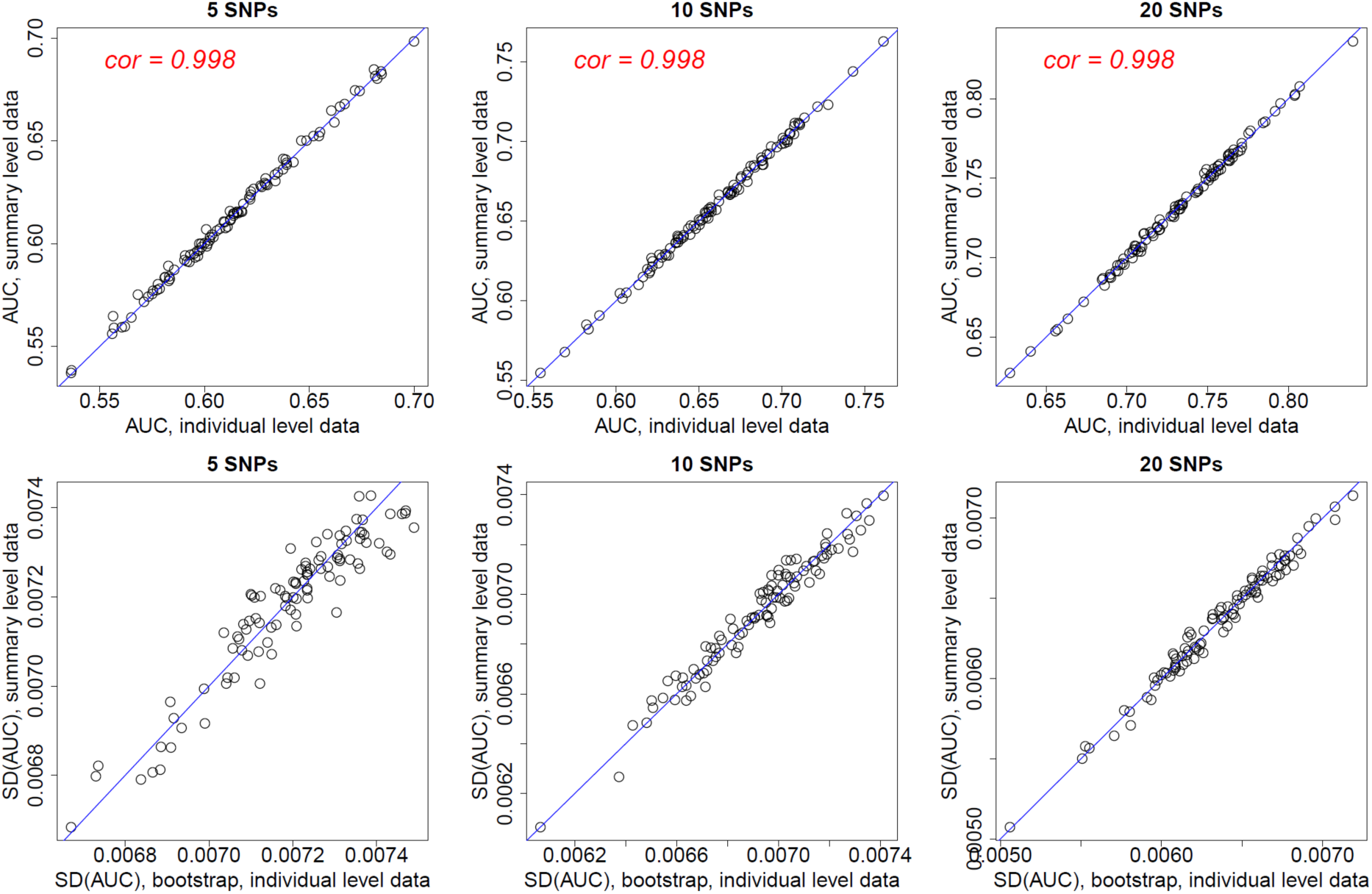
AUC values and the standard errors for PRS with independent SNPs based on simulation study. Each data point represents one simulation. For each simulation, we calculated the AUC and its variance based on individual data (x-coordinate) and using SummaryAUC (y-coordinate).

We then performed simulations by simulating 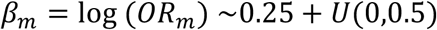. The minimum *OR* is exp(0.25) = 1.28 and the maximum OR is exp(0.75) = 2.11. These values are quite big for common SNPs and polygenic diseases. Results are in Supplemental Figures 1. Again, SummaryAUC provides good approximations.

### 3.3 Application to genetic risk prediction of Schizophrenia

We evaluated the performance of SummaryAUC using schizophrenia GWAS(Schizophrenia Working Group of the Psychiatric Genomics, 2014). Schizophrenia is a devastating psychiatric disorder with high heritability (80-85%) and has a prevalence of approximately 1% worldwide. Schizophrenia is highly polygenic and estimated to be caused by more than 10,000 common SNPs. The Schizophrenia Working Group of Psychiatric Genetics Consortium (PGC) recently performed a meta-analysis of 49 case-control studies (34,241 cases and 45,604 controls) and 3 family studies (1,235 parent affected offspring trios) and identified 108 genome-wide significant common SNPs. Encouragingly, PRSs have achieved a high discrimination performance with average AUC ~75% by leave-one-out analysis. Thus, this data set is very useful for evaluating the performance of SummaryAUC because we can choose PRS with wide range of AUC values. Another advantage of using schizophrenia GWAS data is that the performance is typically maximized when most of SNPs are included in the PRS (Schizophrenia Working Group of the Psychiatric Genomics, 2014), i.e., the p-value threshold for including SNPs in PRS is nearly one. Thus, we can evaluate the accuracy of our method in presence of extensive correlations between SNPs.

PRSs were constructed using the results of a fixed-effect meta-analysis using all sub studies excluding the Molecular Schizophrenia Genetics (MGS) study (Shi, et al., 2009). The MGS study (2681 cases and 2653 controls of European ancestry) were used to calculate AUC values as the independent validation data set. The meta-analysis results included *p*-values *P*_*m*_ and odds ratios *OR_m_* based in single variant logistic regression analysis. Let *w_m_* = log(*OR_m_*).

PRSs were built in two steps following Purcell’s approach (Purcell, et al., 2009): (1) LD-clumping using pairwise correlation threshold *r*. LD-clumping was guided by the association p-values in the training data set to keep the SNPs with smaller p-values in each specified short interval. After LD-clumping, reminding SNPs (denoted as *A_R_*) had pairwise correlation less than r.(2) The PRS was defined as 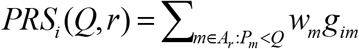 for a given p-value threshold *Q*.

Note that the number of SNPs in PRS increases with *Q* and |*r*|. We chose a wide range of p-value threshold, ranging from 5×10^−8^ (genome-wide significance) to 1 (i.e., all SNPs in *A_r_*). To investigate how the pairwise correlation in SNPs impacted the performance, we chose *r*^2^ = 0.01 (very stringent, SNPs practically independent), 0.1 and 0.2 (locally, modestly correlated).

#### Compare performance when SNPs are genotyped in MGS

In the first set of analyses, we compared AUCs and SEs for PRSs restricted to genotyped SNPs in the MGS dataset. For each PRS, we calculated AUC and its standard error (SE) using MGS as the validation data set in two ways. First, we assumed that individual-level genotype/phenotype data in MGS were available, calculated PRS for each subject in MGS and calculated AUC using an R package “AUC”. Then, we performed bootstrap (N=10000) to estimate the SEs of the estimated AUC values. We denote the two values as AUC_0_ and SE_0_. Second, we performed single variant logistic regression to derive p-values, allele frequencies in controls and odds ratios for all common SNPs, adjusting for sex, age and the top 10 PCA scores. The AUC and its SE were calculated using SummaryAUC. We denote the two values as AUC_1_ and SE_1_.

Results are reported in Figure 3. As was reported previously (Purcell, et al., 2009; Schizophrenia Working Group of the Psychiatric Genomics, 2014), including more SNPs in PRS increases AUC for schizophrenia because of the extremely highly polygenic genetic architecture.

**Figure 3.**
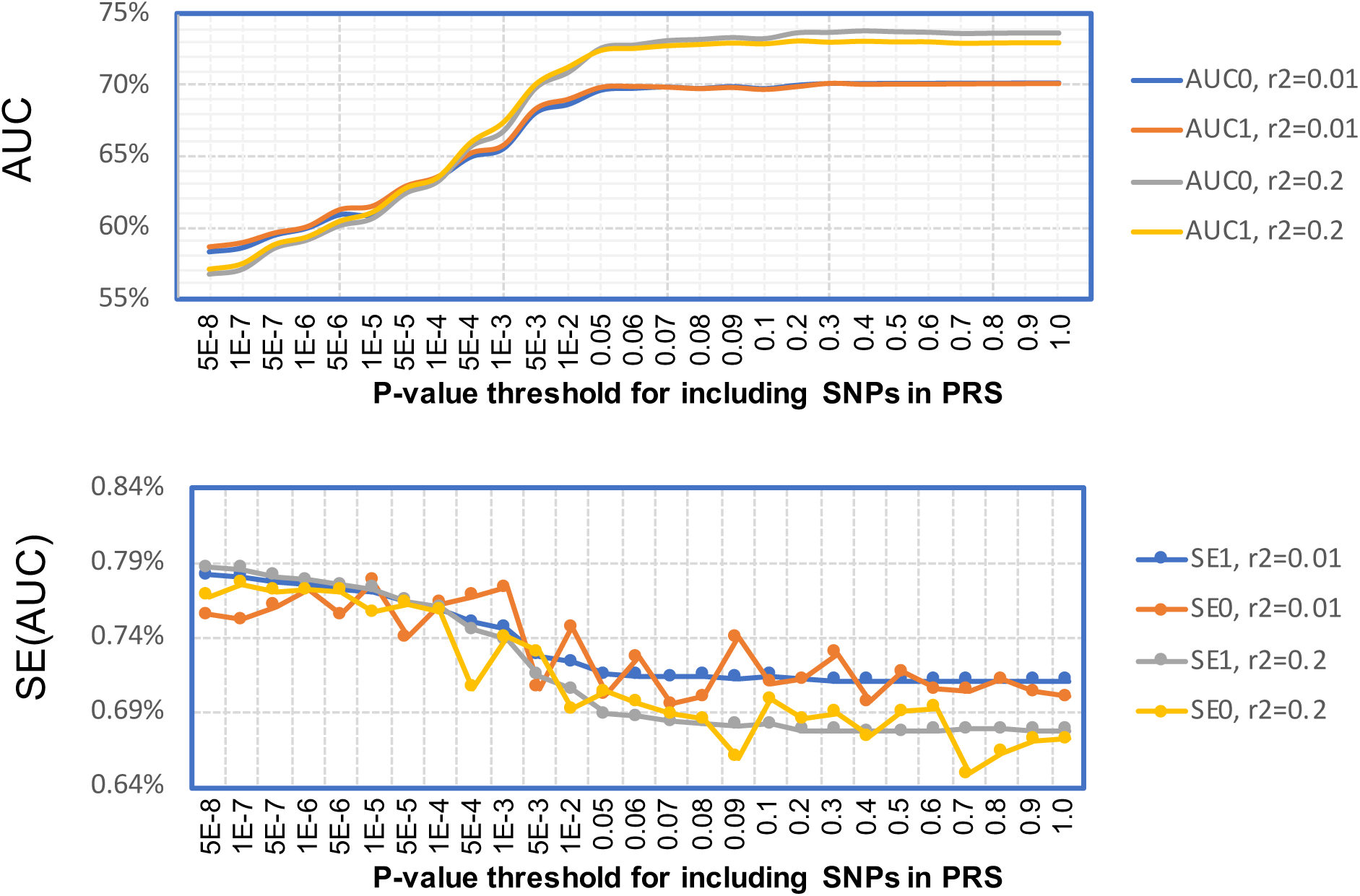
AUC values and their standard errors for PRS of schizophrenia for genotyped SNPs in the MGS study. PRSs were trained based on the PGC summary data excluding the MGS study; the MGS study was used as the validation data for calculating AUC and its variance. Analysis was restricted to SNPs genotyped in MGS. The x-coordinate is the *p*-value threshold (for training data set) for including SNPs in PRS. AUC0 and SE0 were calculated using individual level data. AUC1 and SE1 were calculated using SummaryAUC based on GWAS summary data in MGS. *r*^2^ = 0.01: SNPs were LD-clumped using *r*^2^ = 0.01 in PLINK. *r*^2^ = 0.2: SNPs were LD-clumped using *r*^2^ = 0.2 in PLINK.

When SNPs are practically independent with pruning criteria *r*^2^=0.01, AUC_0_ and AUC_1_ agree very well with the largest difference |AUC_0_-AUC_1_|=0.37%. When we allow SNPs to be weakly correlated with pruning criteria *r*^2^ = 0.2, we observed highly concordant results until *p*-value threshold <0.1, where the largest difference |AUC_0_-AUC_1_|=0.31%. When we included SNPs with more liberal *p*-values, we observed larger inconsistency but the difference is still acceptable with the largest difference is |AUC_0_-AUC_1_|=0.63% when all SNPs (122,552 SNPs) after pruning are included in PRS. Because we only ran 10,000 bootstrap samples to derive SE_0_, there is some fluctuation across PRS models. Apparently, SE_1_ provides an accurate approximation to SE_0_.

To empirically examine the accuracy for PRSs with a small number of SNPs, we examined the performance of SummaryAUC for PRS with the number of SNPs varying from 5 to 100 (Figure 4). When *r*^2^ = 0.01, the largest difference |AUC_0_-AUC_1_|=0.41%; When *r^2^* = 0.2, the largest difference |AUC_0_-AUC_1_|=0.55%. SE values also agree well although SE_0_ fluctuates because of the limited number of bootstrap samples.

**Figure 4.**
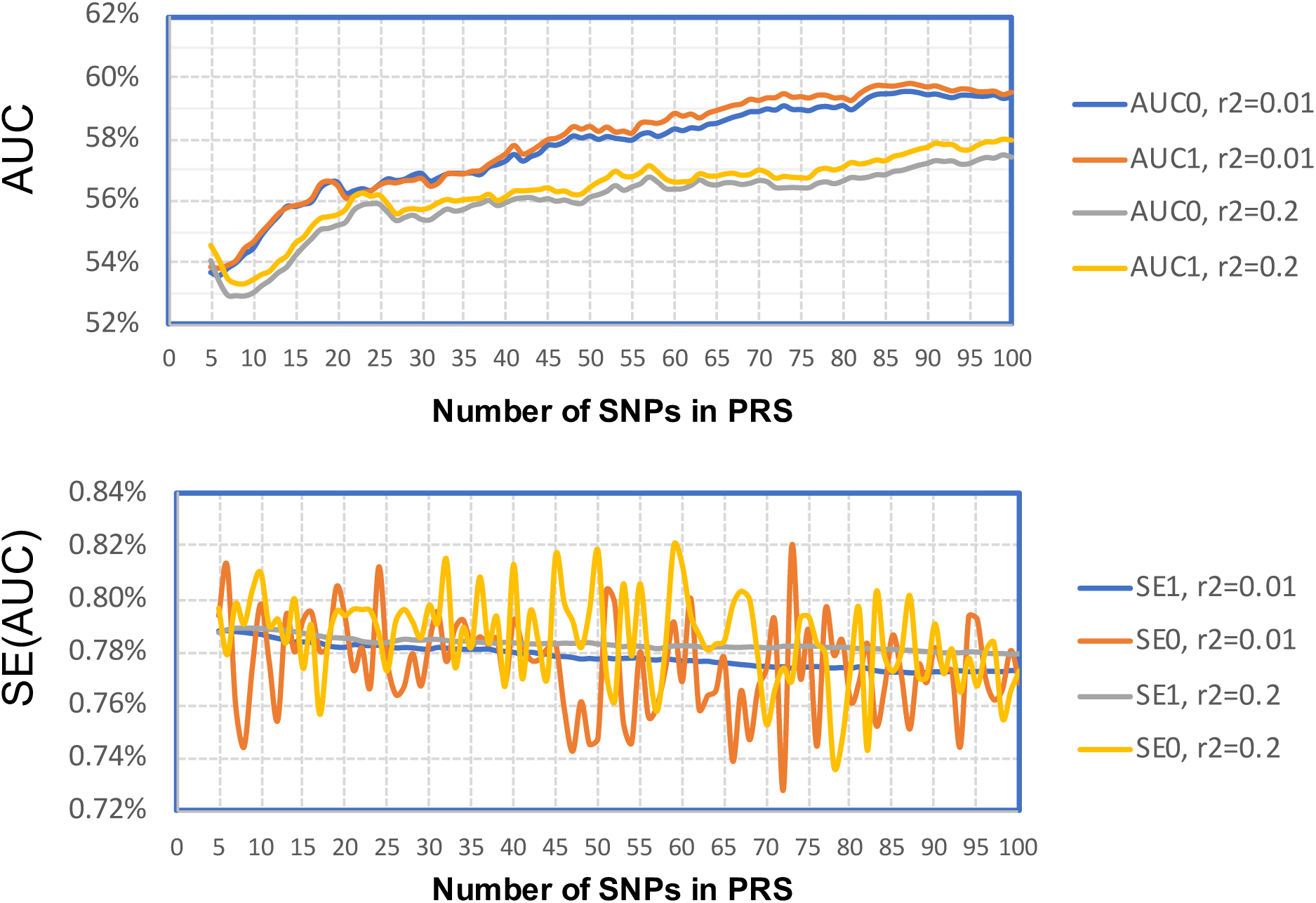
AUC values and their standard errors for PRS of schizophrenia for genotyped SNPs in validation dataset. PRSs were trained based on the PGC data excluding the MGS study; the MGS study was used as the validation data for calculating AUC and its variance. The x-coordinate is the number of SNPs (with smallest p-values in the training data set) for PRS. AUC0 and SE0 were calculated assuming individual level data. AUC1 and SE1 were calculated using SummaryAUC based on GWAS summary data in MGS. *r*^2^ = 0.01: SNPs were LD-clumped using *r*^2^ = 0.01 in PLINK, *r*^2^ = 0.2: SNPs were LD-clumped using *r*^2^ = 0.2 in PLINK.

#### Compare performance when SNPs are imputed in MGS

MGS samples were imputed using software IMPUTE2 (Howie, et al., 2009) and using the haplotypes in The 1000 Genome Project as the reference. SNPs with imputation *R*^2^ < 0.5 were excluded from analyses. Again, AUC_0_ and SE_0_ were calculated using individual level data. AUC1 and SE1 were calculated using SummaryAUC with *p_ml_* = *cor*(*g_im_, g_il_*) estimated directly using the genotype data in The 1000 Genome Project. AUC2 and SE2 were calculated using SummaryAUC with 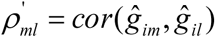 estimated using the imputed genotypic dosage data for samples in The 1000 Genome Project, as illustrated in Figure 1.

Results are presented in Figure 5. When SNPs were very rigorously pruned with *r^2^* = 0.01, both methods approximated AUCs and their standard errors very well. When SNPs were pruned with *r^2^* = 0.2, both AUC1 and AUC_2_ approximate AUC_0_ very well when the p-value threshold <0.01. In fact, when p-value threshold = 0.01, the PRS has 27,404 SNPs. When PRSs use more liberal p-value threshold to increase the number of SNPs, precisely adjusting for local correlation between SNPs becomes more important but difficult for SummaryAUC. In this case, AUC_1_ does not approximate AUC_0_ very well with the largest bias |AUC_0_-AUC_1_|=2.25%. This is because AUC_1_ uses *p_ml_* = *cor* (*g_im_, g_il_*) estimated directly using the genotype data in The 1000 Genome Project. In fact, imputation may change the correlation for imputed SNPs, particularly for poorly imputed SNPs. Encouragingly, AUC_2_ better approximates AUC_0_ with the largest bias |AUC_0_-AUC_2_|=1.30%. The remaining bias may be due to the subtle difference of LD between the external genotype and the MGS population. A similar pattern is observed for SE_0_, SE_1_ and SE_2_.

**Figure 5.**
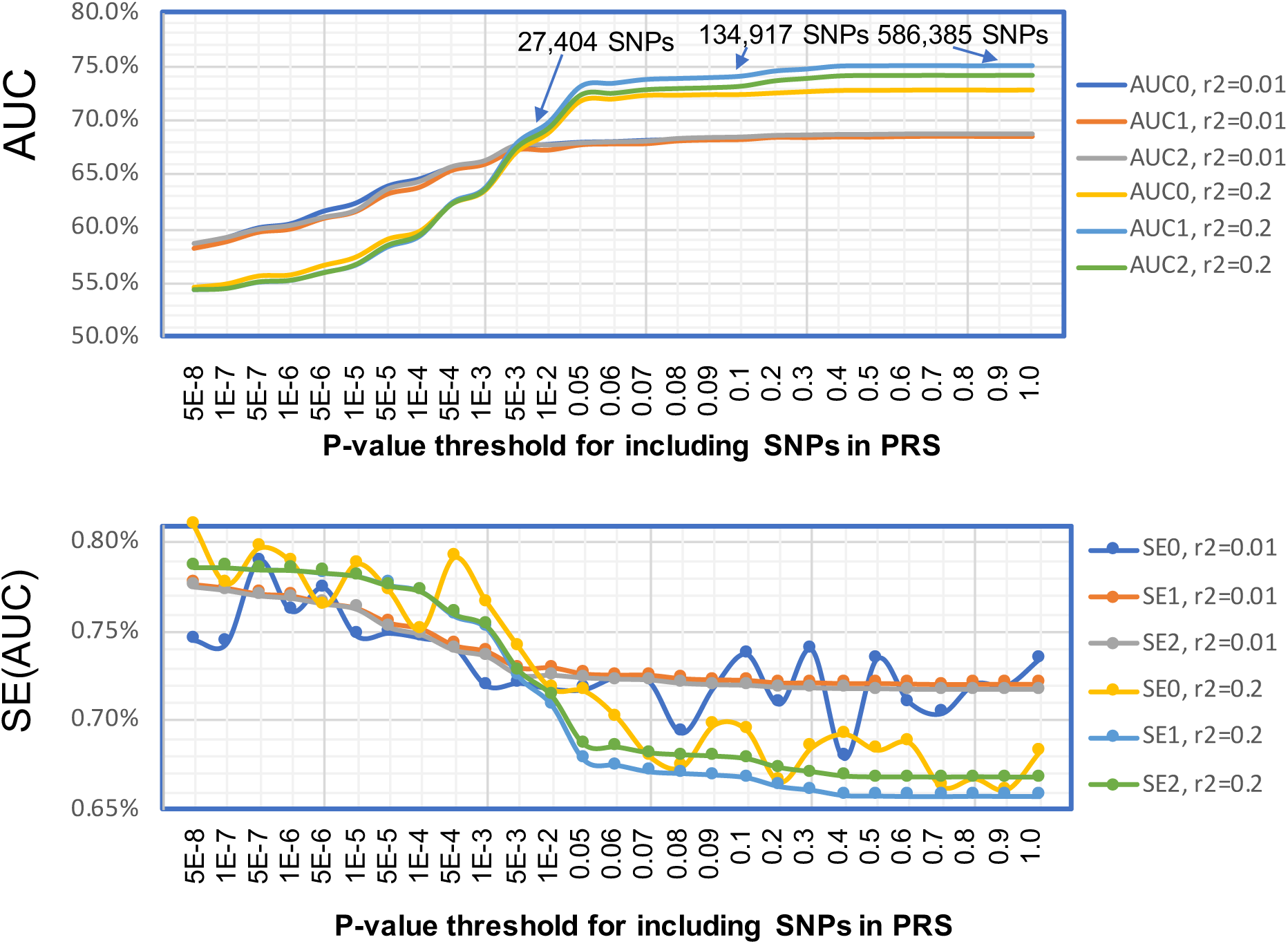
AUC values and their standard errors for PRS of schizophrenia when SNPs are imputed. PRSs were trained based on the PGC data excluding the MGS study; the MGS study was used as the validation data for calculating AUC and its variance. MGS samples were imputed using IMPUTE2 and the haplotypes in The 1000 Genome Project. AUC0 and SE0 were calculated assuming individual level data. AUC1 and SE1 were calculated using SummaryAUC based on GWAS summary data in MGS, where pairwise SNP correlation was estimated using the genotype in The 1000 Genome Project directly. AUC2 and SD2 were calculated using SummaryAUC based on the summary data in MGS, where pairwise correlation was estimated using the imputed genotypic dosage data illustrated in Figure 1. *r*^2^ = 0.01: SNPs were LD-clumped using *r*^2^ = 0.01 in PLINK. *r*^2^ = 0.2: SNPs were LD-clumped using *r*^2^ = 0.2 in PLINK.

## 4. DISCUSSIONS

Although the predictive performance of polygenic risk score (PRS) models are relatively poor for most of complex diseases, PRS will be improved by increasing the sample size of the training GWAS data set and innovative statistical methods that incorporate additional biological information, e.g., functional annotation data and genetic pleiotropy (Chen, 2018; Hu, et al., 2017; Shi, et al., 2016). One difficulty for developing more accurate PRS is to evaluate the predictive performance of PRS in independent GWAS, which requires individual level genotypic and phenotypic data. Herein, we develop SummaryAUC as a new statistical method for approximating AUC and its standard error based on GWAS summary level data, which will greatly facilitate the development of more accurate PRS models.

Although SummaryAUC was derived under the normality assumption of PRS, simulation results suggest that the performance of SummaryAUC is robust to the number of SNPs in PRS. In fact, we found that SummaryAUC was very accurate even for PRS with only five SNPs.

We systematically examined the accuracy of SummaryAUC by applying it to schizophrenia GWAS. Because of the extremely polygenic genetic architecture and the very large sample size I the training dataset, PRS can reach as high as 75% when all SNPs after LD-pruning are included in PRS. This data is very useful for examining the performance of SummaryAUC because we can compare accuracy for a wide range of AUC and for PRSs with tens of thousands of SNPs. The observations from this numerical study can be summarized as follows.

First, when SNPs in PRS are practically independent after rigorous LD pruning, SummaryAUC is most accurate and the accuracy is not compromised even when all SNPs (after pruning) are included in PRS. In this case, computation is very fast.

Second, if locally correlated, genotyped SNPs are included in PRS, the performance is only slightly compromised compared to that based on independent SNPs. Because we have to adjust for local correlation, the computation might be slightly slow. In our implementation, we set *ρ_ml_* = 0 for SNPs on different chromosomes or on the same chromosome but 5 Mb away; thus, computation can be done within a few minutes even when PRS has tens of thousands of SNPs.

Third, it is more complicated when PRS has correlated SNPs that are imputed in the validation GWAS dataset. When PRS is reasonably sparse (e.g., < 20,000 SNPs), SummaryAUC is still accurate. However, when PRS has many SNPs (e.g., when including SNPs using p-value threshold 0.01), SummaryAUC is less accurate if pairwise correlations are not appropriately adjusted. In this case, an imputation-based method helps to reduce the bias. In reality, this only applies to schizophrenia and a few other psychiatric disorders because of their highly polygenic genetic architecture. For most of other diseases, the PRS with optimal classification accuracy is sparse, typically with less than 2000 SNPs. Thus, we expect SummaryAUC to work well.

Thus, if PRSs use independent SNPs, SummaryAUC can be used with confidence for PRS with any size and for both genotyped and imputed SNPs. If PRSs use correlated SNPs that are imputed in validation GWAS dataset, SummaryAUC is most accurate for sparse PRS models and needs to adjust correlations using imputed dosage data only for very dense PRS models. In addition, SummaryAUC is not suitable for PRSs using all common SNPs in the genome, e.g., LD-Pred (Vilhjalmsson, et al., 2015) and BLUP-type PRSs (Golan and Rosset, 2014; Speed and Balding, 2014) that are based on linear mixed models. It is computationally infeasible to adjust for the correlation for millions of SNPs.

Currently, we are working on developing methods for approximating *R*^2^ = *cor*^2^(*y_i_, PRS_i_*)using GWAS summary data in the validation dataset, the fraction of phenotypic variance explained by a PRS model at the observational scale. In addition, we are working on developing statistical methods for testing whether the AUC values from two PRS models are statistically different using GWAS summary data.

### APPENDIX: The variance of AUC estimator based on summary statistics

We calculate the variance of the AUC estimator 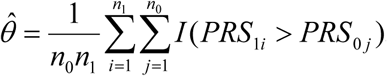. Note that

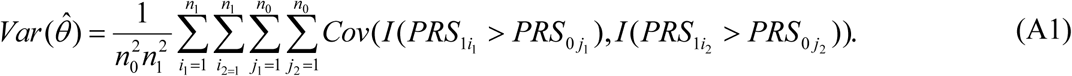

Some algebra shows that (A1) can be partitioned into the four parts:

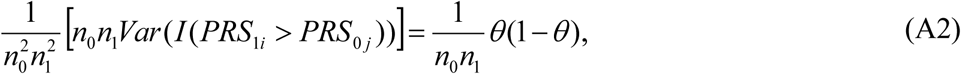

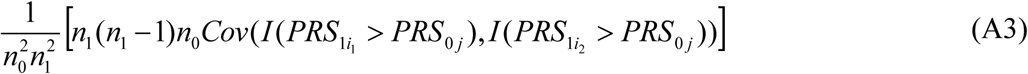

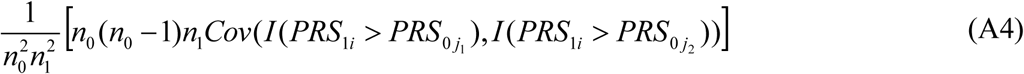

and

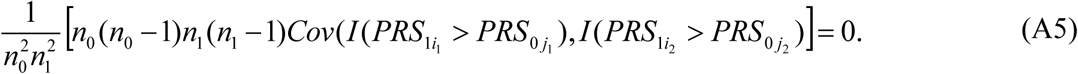

Furthermore, (A3) can be simplified as:

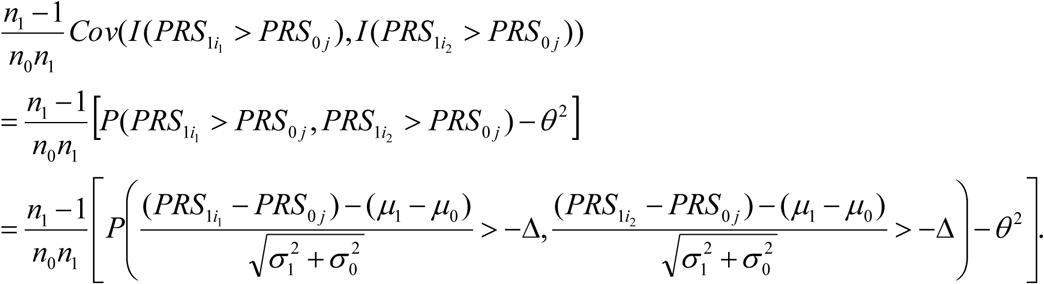

Let 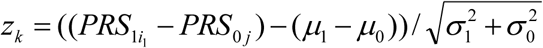 for *k* = 1 and 2. It is easy to verify that 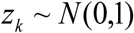 and 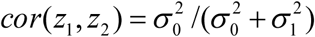.Thus, (A3) is simplified as

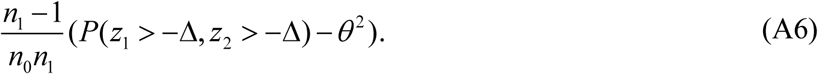

Similarly, (A4) can be simplified as

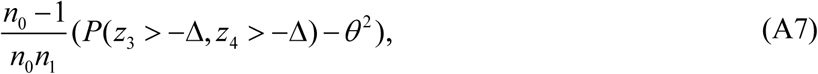

where 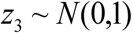, 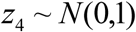 and 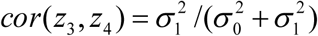

Combining (A1), (A2), (A5), (A6) and (A7) leads to

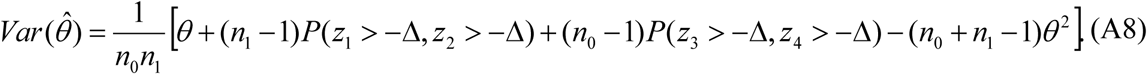

Note that 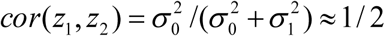 and 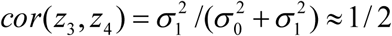 assuming 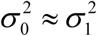. This assumption together with the fact that 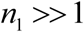 and 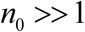 simplifies 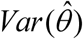 as

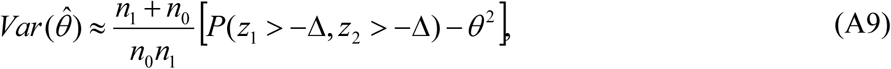

where 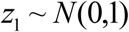, 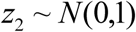 and *cor*(*z*_1_, *z*_2_) = 1/2.

## ACKNOWLEDGEMENTS

This study utilized the high-performance computational capabilities of the Biowulf Linux cluster at the National Institutes of Health, Bethesda, MD. (http://biowulf.nih.gov). The project was supported by the NIH Intramural Research program. The Molecular Genetics of Schizophrenia (MGS) Consortium includes P.V. Gejman, A.R. Sanders, J. Duan (North Shore University Health System and University of Chicago), C.R. Cloninger, D.M. Svrakic (Washington University, St. Louis), N.G. Buccola (Louisiana State University Health Sciences Center, New Orleans), D.F. Levinson, J. Shi (Stanford University, Stanford, Calif.; Dr. Shi is now at the National Cancer Institute), B.J. Mowry (Queensland Centre for Mental Health Research, Brisbane, and Queensland Brain Institute, University of Queensland, Brisbane), R. Freedman, A. Olincy (University of Colorado Denver), F. Amin (Atlanta Veterans Affairs Medical Center and Emory University, Atlanta), D.W. Black (University of Iowa Carver College of Medicine, Iowa City), J.M. Silverman (Mount Sinai School of Medicine, New York), and W.F. Byerley (University of California, San Francisco).

Conflict of Interest: none declared.

